# Spatial Transcriptomic Analysis Identifies a *SERPINA3*-Expressing Astrocytic State Associated with the Human Neuritic Plaque Microenvironment

**DOI:** 10.1101/2024.11.13.623438

**Authors:** Berke Karaahmet, Ya Zhang, Laurine Duquesne, Alina Sigalov, Christina Yung, Alexandra Kroshilina, David A. Bennett, Mariko Taga, Hans-Ulrich Klein

**Affiliations:** Center for Translational & Computational Neuroimmunology, Department of Neurology and Taub Institute for Research on Alzheimer’s Disease and the Aging Brain, Columbia University Medical Center, New York, New York, USA; Rush Alzheimer’s Disease Center, Rush University Medical Center, Chicago, IL, USA

**Keywords:** Spatial Transcriptomics, Alzheimer’s Disease, Astrocytes, SERPINA3

## Abstract

Single-nucleus transcriptomic studies have revealed glial cell states associated with Alzheimer’s disease; however, these nuclei are dissociated from the complex architecture of the human neocortex. Here, we successfully performed an unbiased distance-based analytic strategy on spatially-registered transcriptomic data. Leveraging immunohistochemistry in the same tissue section, our analyses prioritized *SERPINA3* and other genes, such as metallothioneins, as altered in the vicinity of neuritic amyloid plaques. Results were validated at the protein level by immunofluorescence, highlighting that a reactive SERPINA3^+^ astrocyte subtype, Ast.5, plays a role in the plaque microenvironment.

## Main

Amyloid-beta (Aβ) deposits in the form of neuritic plaques are a pathological hallmark of Alzheimer’s disease (AD)^1^. Recent single-nucleus transcriptomic characterizations of glial cell populations in AD brains have revealed associations between pathological AD traits and cognitive decline with the emergence of specific microglial and astrocytic states^2-5^. However, the complex multicellular processes involved in plaque formation or in response to plaques *in situ* remain poorly understood due to the loss of histopathological spatial information. In this study, we sought to characterize the transcriptomic landscape within the neuritic plaque microenvironment using the 10x Genomics Visium Spatial Transcriptomics (ST) platform^6^.

Previous studies have utilized ST to examine the inflammatory response to Aβ pathology in mouse models^7,8^ and have validated these findings in human brains using targeted assays^9^. However, these approaches may overlook changes specific to humans. Furthermore, earlier research combined ST methods with immunohistochemistry (IHC) on adjacent tissue sections to identify differential expression around Aβ plaques^8,9^. In contrast, our study employed an adjusted IHC protocol compatible with Visium ST library preparation on the same fresh-frozen human brain tissue sections (Figure 1a). This method reduces errors introduced by tissue preparation, image registration, and plaque projection, thus providing a more accurate assessment of transcriptional changes in the neuritic plaque microenvironment.

**Figure 1.**
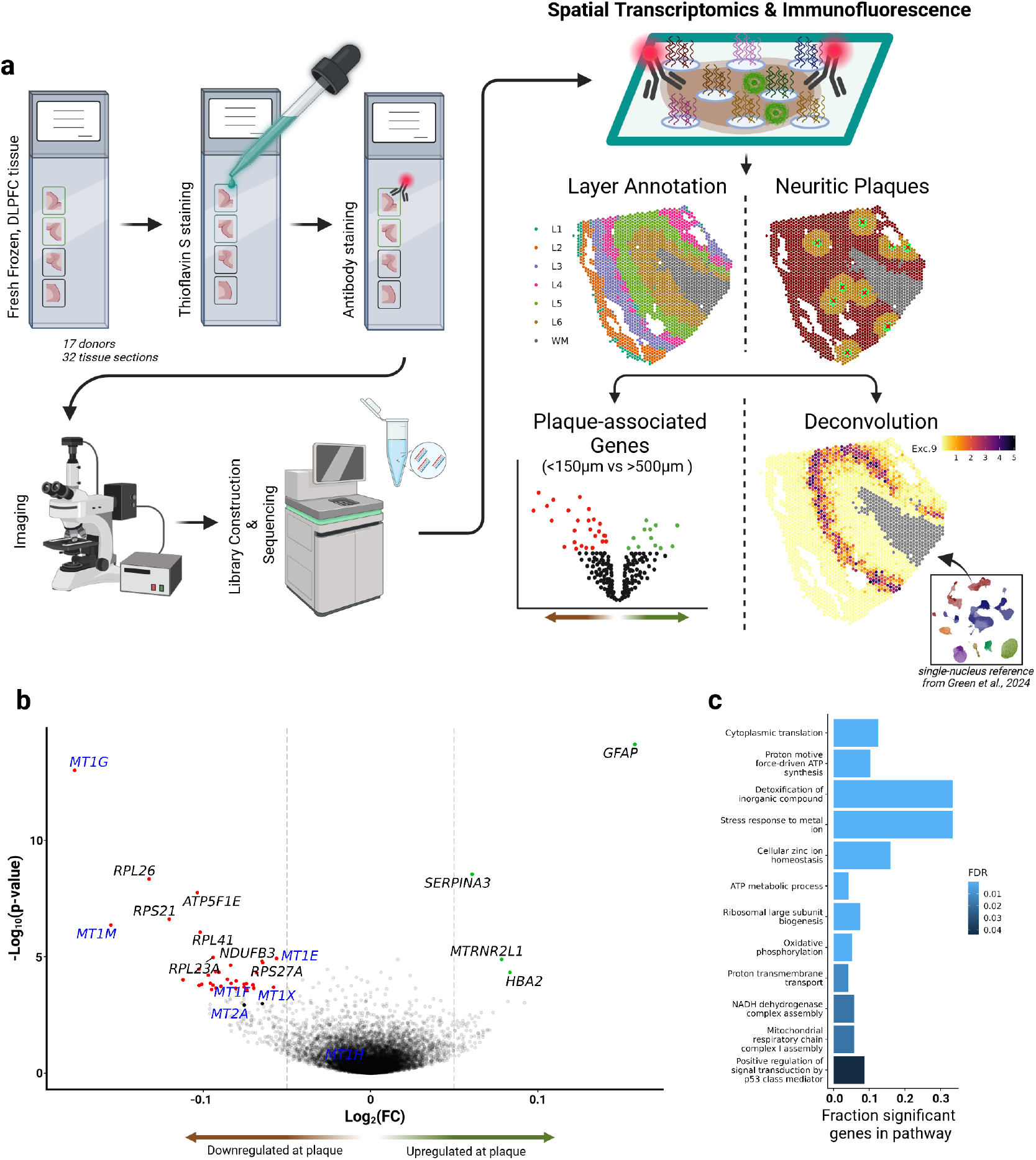
**(a)** Schematic of experimental workflow. Fresh-frozen DLPFC tissue from 17 donors were loaded in duplicates on 10x Genomics Visium slides. The tissues were stained with ThioS and anti-GFAP using the modified IHC protocol as described in *Methods*. Imaging was performed immediately after the staining followed by library construction and sequencing. The locations of neuritic plaques were manually identified using ImageJ software. Neuronal layers, which contribute the most to variability in this ST dataset, were annotated using spatially-aware clustering with BayesSpace. We then conducted differential expression testing between ST spots within 150μm of neuritic plaques compared to spots greater than 500μm away from neuritic plaques, correcting for layer and donor information. In parallel, we performed computational deconvolution of the ST data using our snRNA-seq dataset that included donor-matched nuclei isolated from DLPFC. Estimated cellular abundance of excitatory neuronal state 9 (Exc.9) is shown here as an example. We used the deconvolution results to identify glial states enriched within 150μm of neuritic plaques. **(b)** Volcano plot shows the 4 upregulated (green) and 35 downregulated (red) genes in spots proximal to neuritic plaques. Two additional metallothionein genes *MT1X* and *MT2A*, colored in black, reached nominal significance. All the metallothionein genes detected in this dataset are labeled in blue. *GFAP* and *SERPINA3* are the most significantly upregulated genes. **(c)** Gene ontology analyses revealed a dysregulation of pathways involved in ion homeostasis. This signature is largely driven by the downregulation of metallothionein genes, as 6 out of 7 detected metallothioneins were downregulated (log_2_ Fold Change < -0.05).

Here, we aimed to identify genes whose expression levels were related to their physical proximity to a key pathologic feature of AD, the neuritic amyloid plaque. We generated ST profiles of 32 fresh-frozen tissue sections from the dorsolateral prefrontal cortex (DLPFC) of 17 donors, 15 of whom had a clinical AD diagnosis and advanced AD pathology (i.e., Braak Stages 4-5), and 2 cognitively non-impaired donors without a pathologic AD diagnosis. Demographic and clinicopathologic characteristics are presented in Supplementary Table 1. We used the Visium platform: each tissue section is placed on top of a glass slide on which there is a printed array of spatially barcoded collection of capture probes. Each spot in the array has a 55μm diameter. Following RNA capture, IHC for GFAP (as an astrocyte marker), DAPI (for nuclei) and Thioflavin S (ThioS; to detect neuritic plaques) were performed on the same tissue section. Each tissue section was scanned at 10X, and we manually annotated the location of ThioS^+^ neuritic plaques (see Methods).

Following preprocessing and quality control, our dataset consisted of 59,588 tissue-covered spots, with each spot capturing RNA from an average of 3.7 nuclei and detecting an average of 2,361 genes (Supplementary Figure 1a-d). Using our combined IHC and ST approach, we were able to identify the location of 263 ThioS^+^ neuritic plaques among our 32 tissue sections. To assess the validity of this approach, we also quantified GFAP protein levels from the IHC images and GFAP mRNA levels from the ST profiles for each spot per section. This analysis revealed significant correlations between the two data modalities, suggesting that immunofluorescent staining can be reliably combined with the Visium ST protocol (Supplementary Figure 2a-d).

The six cortical layers with their distinct cell types were the main source of variation at the spot level in our ST data and needed to be statistically accounted for in the subsequent analyses. We applied a spatially-aware clustering method using highly variable genes and layer marker genes (see Methods) to detect white matter spots and annotate grey matter spots with cortical layers^10^. Comparing layer-specific transcriptional signatures derived from our dataset with those derived from a published DLPFC ST dataset with H&E-based manual layer annotation showed a strong correlation, confirming the concordance of our clustering-based annotation with classical histopathological annotation (Supplementary Figure 3a)^11^. Furthermore, using our layer annotations and plaque locations, we found a higher rate of plaques in the deeper layers (L4 - L6) compared to the upper layers (L1 - L3), (Supplementary Table 2; Supplementary Figure 3b-e), similar to previously published results^12^.

To perform our primary analysis, we calculated the distance from each spot to the closest neuritic plaque and performed differential expression testing comparing spots within 150μm of a neuritic plaque versus those more than 500μm away, adjusted for cortical layer and donor. This revealed a statistically significant upregulation of *GFAP, SERPINA3, MTRNR2L1*, and *HBA2* at the plaque niche (Figure 1b and Supplementary Table 3). The significantly downregulated genes included various metallothioneins, mitochondrial, and ribosomal genes. Gene ontology analysis revealed a positive enrichment of pathways related to metal ion response, suggesting dysregulation of metal ion homeostasis within the plaque niche (Figure 1c). We next utilized single-nucleus RNA sequencing (snRNA-seq) data previously generated from the same cohort and brain region to explore the cell types expressing these plaque-associated genes. Metallothioneins showed the highest transcription levels in the nuclei of astrocytes, although they were also transcribed by other cell types at lower levels (Supplementary Figure 4a). The upregulated gene *SERPINA3* was almost exclusively transcribed by astrocytes in our human gray matter DLPFC data (Supplementary Figure 4a).

We next sought to confirm the overexpression of SERPINA3 in astrocytes proximal to neuritic plaques at the protein level using an independent set of 20 ROSMAP paraffin-embedded DLPFC tissue sections from donors with Braak stages 5 or 6 (Supplementary Table 1). IHC was performed for Aβ_1-42_, SERPINA3, and GFAP, a classic astrocyte marker and also the most significantly upregulated gene in relation to its distance from neuritic plaques in our analysis. We then segmented DAPI^+^GFAP^+^ astrocytes in the tissue images and quantified the median SERPINA3 and GFAP intensities for each astrocyte (Figure 2a). Comparing astrocytes proximal to neuritic plaques (≤100µm; 12,763 astrocytes) with astrocytes distant to plaques (between 350µm and 500µm; 51,990 astrocytes) revealed a 3.2% increase in median SERPINA3 intensity (p<1×10^−16^) and an 11.8% increase in median GFAP intensity (p<1×10^−16^) in astrocytes at plaques (Figure 2b); these effect sizes are similar to those observed at the RNA level. Compared to the Visium ST analyses, the higher resolution of the IHC images allowed us to use a smaller area around neuritic plaques. We further chose to restrict the control area to less than 500µm as we were unable to adjust for cortical layers in the IHC analysis. Plotting the SERPINA3 and GFAP median fluorescence intensities (MFIs) per astrocyte depending on their distance to neuritic plaques revealed that their levels were highest within 50µm and plateaued around 300µm away from the plaques (Figure 2c).

**Figure 2.**
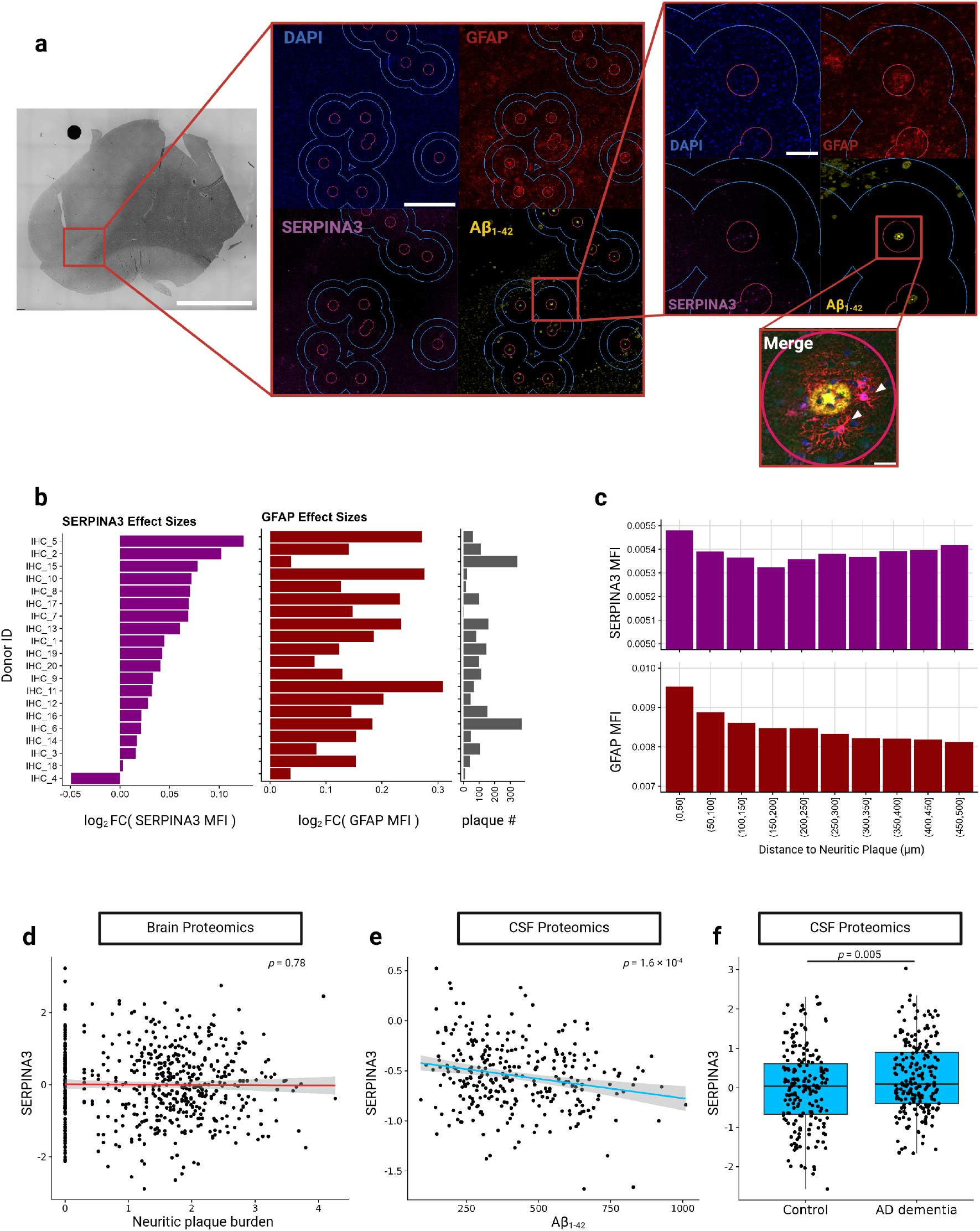
IHC validation of DLPFC tissue from a separate cohort of AD individuals reveals enrichment of SERPINA3 expression by astrocytes within the neuritic plaque microenvironment. **(a)** Representative images and insets showing the analytical approach and SERPINA3^+^ astrocytes. In the zoomed in fluorescent images, astrocytes falling within the 350μm and 500μm of distance from neuritic plaques (i.e. the area between the blue circles) were annotated as being distant from a plaque while astrocytes within 100μm of neuritic plaques (i.e. the area within the pink circles) were annotated as being proximal to plaques. DAPI is shown in blue, GFAP in red, SERPINA3 in purple, and Aβ_1-42_ in yellow. Representatives of SERPINA3^+^ astrocytes are marked with white arrowheads in the merged image. Scale bars denote 5000μm, 1000μm, 200μm and 10μm in their respective magnification order. **(b)** Purple bars depict the log_2_ fold change of SERPINA3 MFI in astrocytes proximal to plaques compared to astrocytes distant from plaques for each of the 20 AD individuals. Red bars indicate the MFI log_2_ fold changes observed for GFAP. The number of neuritic plaques for each donor is shown in grey bars. Notably, all the samples show a positive association between MFIs and plaque proximity, except for one individual. **(c)** Plaque distance-binned average MFIs of SERPINA3 and GFAP of segmented astrocytes are shown in histograms. SERPINA3 and GFAP MFIs are highest in astrocytes within 50μm of a neuritic plaque. **(d)** Proteomic analysis of brain parenchyma reveals no correlation between SERPINA3 and neuritic plaque burden (p=0.78) **(e)** but proteomics of CSF shows a significant negative correlation of SERPINA3 levels with increasing CSF Aβ_1-42_ levels (p=1.6×10^−4^) and **(f)** a significant increase of SERPINA3 levels in those with AD dementia (p=0.005).

We next investigated whether the localized upregulation of SERPINA3 at neuritic plaques could be detected in DLPFC tissue homogenates or the cerebrospinal fluid (CSF) of patients with a high burden of amyloid pathology. Although, we did not observe an association between SERPINA3 protein levels in bulk DLPFC proteomic data and neocortical neuritic plaque loads in 588 ROSMAP participants (Figure 2d)^13^, we identified a significant negative association between SERPINA3 and Aβ_1-42_ levels in the CSF of 296 donors from an independent cohort (p=1.6×10^−4^), with a 5% decrease in SERPINA3 per standard deviation (SD) increase in Aβ_1-42_ (Figure 2e)^14^, noting that low Aβ_1-42_ in CSF reflects high cortical Aβ_1-42_ accumulation. This finding was further validated using a second CSF proteomics dataset, which confirmed significantly higher CSF SERPINA3 levels in 209 AD patients with dementia compared to 187 controls (p=0.005, 20% increase in AD, Figure 2f)^15^. Overall, these data provide evidence for increased SERPINA3 protein in the CSF of donors with higher levels of cortical Aβ.

Recent snRNA-seq studies have highlighted the remarkable heterogeneity of human glial cells, with some transcriptionally distinct glial states being linked to AD pathologies^2-5,16-20^. To determine which of these glial cell states are found in proximity to neuritic plaques, we integrated our snRNA-seq reference data^2^, generated from the same brain region and cohort, with our ST data. Using the cell2location algorithm^21^, we then predicted cell state abundances at each spatial location. The predicted abundances of excitatory neuronal subpopulations reflected the cortical laminar structure, consistent with the layer-specific annotations of these subpopulations from the Allen Brain Map (Supplementary Figure 5a-d). This confirmed the accuracy of the cell2location predictions. Among the astrocytic and microglial states, Astrocyte state 5 (Ast.5) was the most significantly enriched state (FDR=6.8×10^−7^, 11.9% higher odds to observe Ast.5) within 150μm of neuritic plaques compared to distant areas (≥500μm) (Figure 3a-d). Consistently, Ast.5 exhibited a reactive-like transcription signature in the snRNA-seq data with high levels of *GFAP* and *SERPINA3* expression (Supplementary Figure 4b), two of the four genes significantly upregulated at neuritic plaques^2,22^.

**Figure 3.**
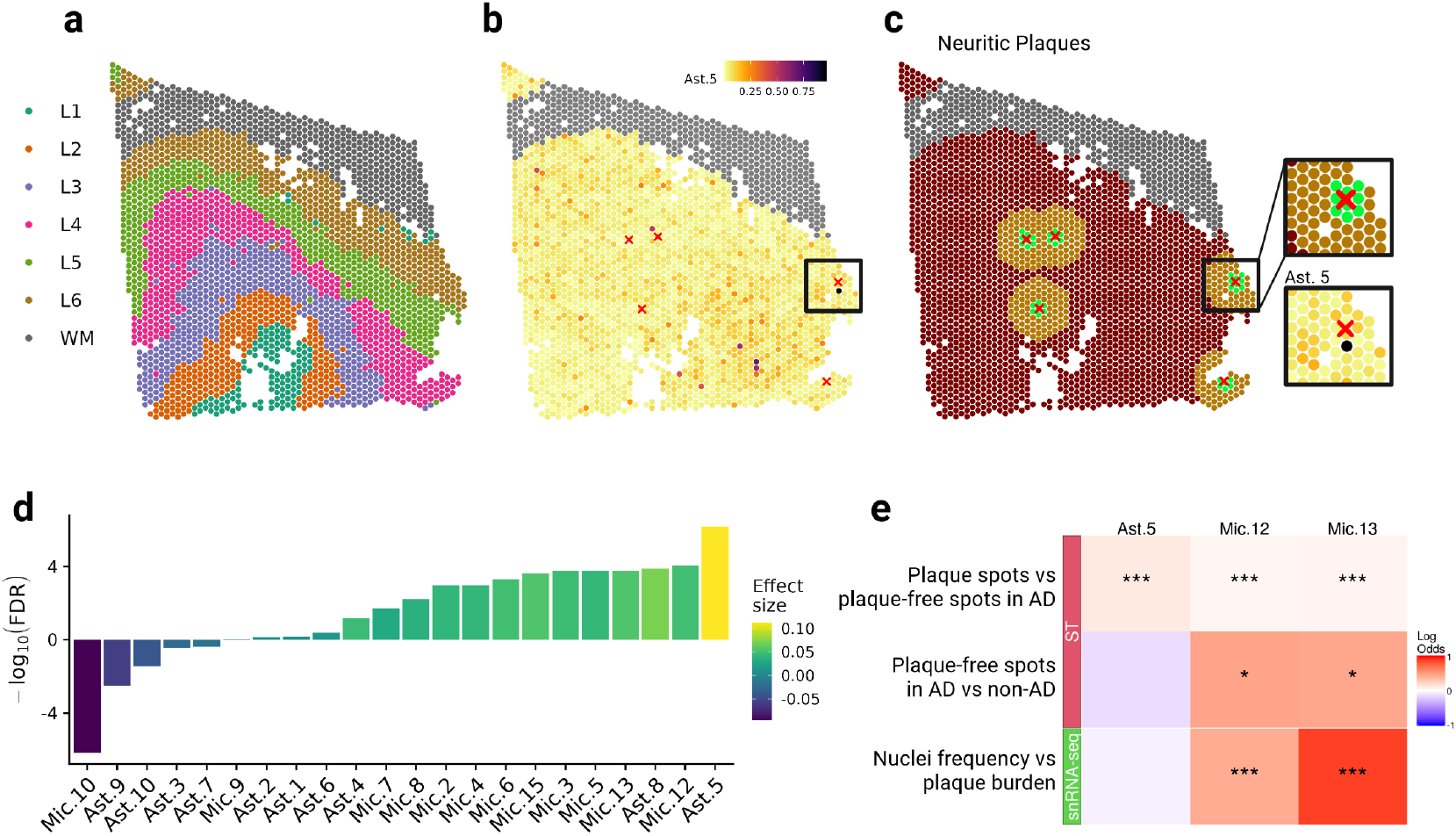
Analysis of transcriptomic signatures identifies positive enrichment of the Ast.5 state at the neuritic plaque microenvironment. **(a)** Representative tissue section denoting the layer structure, **(b)** Ast.5 abundance and **(c)** distance of a given spot from the closest neuritic plaque (green < 150μm, orange < 500μm). White matter is colored grey and neuritic plaques are marked with a red cross (“×”). The inset in **(c)** shows a zoomed in region containing a neuritic plaque (top) and the estimated Ast.5 abundance in the same region (bottom). **(d)** Log odds ratio of estimated microglial and astrocytic state abundances proximal (≤150μm) or distal from neuritic plaques (≥500μm), corrected for layer and donor, reveals that Ast.5 is the most strongly enriched glial state at neuritic plaques. **(e)** Log odds ratios of snRNA-seq data show a global correlation between Mic.12 and Mic.13 frequencies with increased neuritic plaque burden but not for Ast.5 (bottom row). This observation is consistent with log odds ratios of cell2location estimates in non-plaque ST spots between AD and non-AD donor tissues for the same cell states (middle row). On the other hand, Ast.5, Mic.12 and Mic.13 are enriched at neuritic plaques in AD issue in the ST data (top row), suggesting that the increase in Ast.5, unlike Mic.12 and Mic.13, is strictly localized to plaque pathology.

The cell2location analysis also identified microglial cell states enriched at neuritic plaques. The two most significantly enriched microglial states were Mic.12 (FDR=8.6×10^−5^, 5.1% higher odds at plaques) and Mic.13 (FDR=1.8×10^−4^, 5.7% higher odds at plaques). They were characterized by the expression of lipid-associated genes in the snRNA-seq reference data, including high levels of *APOE* and *GPNMB*^2^. Notably, classical disease-associated microglial genes (such as *TREM2, ITGAX, AXL*)^23^ were not among our plaque-associated genes, likely due to the limitations of the Visium ST platform, which does not detect low RNA levels from infrequent cell types reliably, resulting in sparse data for microglia-specific genes. This shortcoming is mitigated by the cell2location deconvolution approach, which utilizes the entire transcriptome. Next, we employed our snRNA-seq reference dataset from 421 donors^2^ to examine whether the frequencies of the plaque-associated cell states, Mic.12, Mic.13, and Ast.5, in the DLPFC correlated with the donors’ neocortical neuritic plaque burden. Interestingly, in the snRNA-seq data, we found strong positive associations for Mic.12 (p<1×10^−16^, 53% higher odds per SD plaque load) and Mic.13 (p<1×10^−16^, 144% higher odds per SD plaque load), but not for Ast.5 (Fig. 3e). Similarly, comparisons of spots distant from plaques (≥500μm) in AD brains with spots from four non-AD sections in our ST data revealed an increase in Mic.12 (p=0.009, 60% higher odds in AD) and Mic.13 (p=0.015, 58% higher odds in AD), but not in Ast.5. Overall, this suggests that Mic.12 and Mic.13 may spread further from plaques, making their association with plaque burden detectable in the dissociated single-nucleus data. The role of these widespread microglial subtypes in relation to neuritic plaques may be fundamentally different from that of the more focal Ast.5.

## Discussion

This study aimed to identify genes and cell subpopulations that are altered in the vicinity of neuritic plaques in the human AD brain. Despite a modest sample size, we found robust results such as *SERPINA3* and *GFAP* (Figure 1b) using our distance-based analysis; results that we validated by immunofluorescence and repurposed CSF data. These results are also consistent with earlier studies^22,24^. Altogether, they provide evidence that our analytic approach is rigorous and robust; scaling it up to a larger sample size will help to uncover a larger array of genes and pathways that have a role in the amyloid proteinopathy which is a key component of AD. Improvements in ST technologies will further enhance analyses as true single-cell spatially-registered data resolves the ambiguity of which cell contributes to which RNA profile.

The plaque-associated astroglial subpopulation, Ast.5, has previously been characterized by high *SERPINA3* expression and it exhibited a transcriptional signature similar to the Disease-Associated Astrocytes (DAAs) that are enriched in the 5xFAD mouse model^22^. Furthermore, human SERPINA3 and its mouse homologues, such as *Serpina3n*, have been shown to interact with plaques in previous studies^7,9,24-27^. While oligodendrocytes in mice also upregulate *Serpina3n* in response to various CNS pathologies^28^; our and others’ snRNA-seq data indicate that astrocytes are the primary source of *SERPINA3* in human gray matter^29^. The increase of Ast.5 may be strictly localized, which could explain why Ast.5 frequencies or SERPINA3 protein levels were not significantly increased in the brain parenchyma in AD. Nevertheless, we found elevated SERPINA3 levels in the CSF of AD patients, confirming earlier studies^30,31^. SERPINA3 in the CSF may be secreted by Ast.5 and could serve as a biomarker for this specific cell state in future studies.

Our analyses also revealed a downregulation of several metallothioneins in regions near neuritic plaques. Metallothioneins are a family of small, cysteine-rich proteins that regulate metal ion homeostasis by chelating or releasing metal ions. Notably, zinc and other metal ions, which are present at high concentrations within plaques, may contribute to plaque formation by binding to Aβ, altering its conformation and solubility, and potentially explaining the accumulation of Aβ oligomers at excitatory synapses, where zinc is released^32-35^. Elevated extracellular zinc levels have been observed in individuals at risk for AD^36^, and the use of metal chelators has been shown to reduce Aβ plaque formation in rodent models^37^. At a cellular level, astrocytes are known to exhibit the highest levels of metallothionein expression in the human brain (Supplementary Figure 4), but other cell types may also contribute. For example, neuronal cell lines treated with Aβ showed a marked downregulation of MT1G *in vitro*^38^. Similarly, microglia from human AD brains demonstrated reduced expression of genes involved in metal homeostasis^29^, whereas microglial cell models stimulated with Aβ showed an upregulation of these genes^39^, suggesting that the observed metal ion dyshomeostasis at plaques arises from a complex, multicellular process.

In summary, we applied a distance-based analytical approach that is anchored to the location of a key pathologic feature of AD. Our analyses prioritized the role of an astrocyte subtype, Ast.5, which did not emerge from the well-powered snRNAseq-based studies, illustrating the limitations of studies that rely solely on changes in the frequencies of cell subtypes. Here, we prioritized a subtype through the rearrangement of its distribution in the affected neocortex. *SERPINA3* is a robust marker of this astrocyte subtype and has been directly implicated in the formation of amyloid plaques^40,41^. Our results present a roadmap for a new analytic strategy to understand the role of selected genes and cell subtypes that have a very specific, topologically restricted role.

## Methods

### Human Postmortem Brain Tissues

The Religious Orders Study and the Rush Memory and Aging Project (ROSMAP) are two cohort studies of aging and dementia^42^. ROSMAP participants enroll without known dementia and undergo annual cognitive tests, and upon death, a detailed neuropathologic examination, including Braak and CERAD staging. The ROS and MAP studies were approved by an Institutional Review Board of Rush University Medical Center. All participants signed an informed consent, an Anatomic Gift Act for brain donation, and a repository consent to allow their data and biospecimens to be shared. Frozen and fixed brain tissues are available to researchers upon request (www.radc.rush.edu). We selected DLPFC tissues from 15 participants with advanced Braak stages (4-5) and CERAD scores of ‘probable AD’ or ‘definite AD’ who also received a clinical AD dementia diagnosis. Additionally, we included 2 participants without clinical or pathologic AD (Braak stages 0 or 1, CERAD score of ‘no AD’). The samples selected for this study were included in a previously published snRNA-seq dataset of 424 ROSMAP DLPFC tissues. The snRNA-seq dataset and the respective cell annotations were used as a reference to deconvolute and interpret our ST data and are described in detail elsewhere^2^.

### Spatial Transcriptomics data generation and analysis

#### Visium data generation

The Visium ST platform and immunofluorescence were applied to tissue sections obtained from fresh-frozen DLPFC samples from ROSMAP decedents. Each section encompassed all cortical layers and some underlying white matter. Only samples with an RNA Integrity Number (RIN) greater than 6 were selected. RNA was purified using the RNeasy Micro Kit (Qiagen, catalog number 74004), and RIN was measured using TapeStation 4150 and Bioanalyzer 2100 (Agilent Genomics).

Tissue blocks were dissected on dry ice to be prepared into approximately 6 × 6mm^2^ sized tissue sections embedded in Optimal Cutting Temperature (OCT; Sakura, 4583) and cut into 10 μm sections in a cryostat and mounted on Tissue Optimization Slides and Gene Expression Slides (10x Genomics) in duplicates.

Permeabilization time was optimized following the manufacturer’s instructions (10x Genomics, Visium Spatial Tissue Optimization, CG000238 Rev D) with modifications. In brief, sections were fixed with cold 100% ethanol (Fisher Scientific, BP2818) for 30 min at –20ºC and stained with Thioflavin S (ThioS; Sigma, T1892) for 7 min at Room Temperature (RT). After blocking with Fc Receptor Blocking Solution (Biolegend, Cat #422301) containing RNAse inhibitor (Thermofisher Scientific, EO0384), the tissues were stained with anti-GFAP Cy3 conjugate (Millipore Sigma, MAB3402C3, dilution 1/50) for 30 min RT. After serial washing with the wash buffer, the slides were incubated with TrueBlack (Biotinum, 23007) prepared in 70% ethanol for 2 min to quench the autofluorescence from lipofuscin. The slides were then immersed in the 3X SSC Buffer (Millipore Sigma, S6639L) for a few seconds at RT and coverslipped with a mounting medium solution containing 85% Glycerol, 2U/µl RNase inhibitor and DAPI solution (Thermofisher Scientific, 62248). The sections were then scanned using the Nikon Eclipse Ni-E immunofluorescence microscope at 10X magnification (Plan Apo λ, NA = 0.45). After imaging, the coverslip was removed and the fixed and stained tissue sections on the Visium Spatial Tissue Optimization Slide were incubated with the Permeabilization Enzyme for different lengths of time.

Three minutes of permeabilization time was chosen for the Visium Spatial Gene Expression workflow to ensure sufficient mRNA release and minimize mRNA diffusion during library preparation. Tissue slices were then processed for gene expression following the manufacturer’s instructions (10x Genomics, Visium Spatial Gene Expression Reagent Kits, CG000239 Rev D). In brief, the stained tissue sections were permeabilized for 3 minutes, followed by reverse transcription, second-strand synthesis, and denaturation. qPCR experiment was processed using KAPA SYBR FAST kit (KAPA Biosystems) and QuantStudio 6 Flex system (ThermoFisher). The cDNA amplification cycle number was determined by ∼25% of the peak fluorescence value. Subsequently, libraries were sequenced on a NovaSeq 6000 system (Illumina), and ST spots (diameter = 55µm) were aligned with the brightfield images using the Loupe Brower (8.0.0, 10x Genomics). ThioS^+^ neuritic plaques were identified by visual inspection by a neuropathology expert. Euclidian distance was used to measure the distance of a given spot to the nearest neuritic plaque.

#### Visium spot-level data processing and cortical layer annotations

Visium sequence reads were aligned to the human reference genome (GRCh38) and assigned to spots using Space Ranger (2.0.0, 10x Genomics). The h5 file output by Space Ranger for each sample was read in R (4.2.2) using the SpatialExperiment package. Sections with a median UMI count less than 2,000, a median detected number of genes less than 1,000, or less than 1,000 spots covered by the tissue section were deemed low quality and removed (4 out of 38 sections). Another 2 sections of primarily white matter were removed, resulting in 32 high-quality sections. Within high-quality sections, single spots with less than 500 detected genes or less than 1,000 UMIs were removed (4.5% of the 62,405 spots).

Spot-level data were normalized by scaling and log-transforming the counts using logNormCounts from the scuttle package (1.4.8)^43^. To detect the cortical layers and the white matter, the top 10,000 highly variable genes were determined using modelGeneVar in the scran package (1.26.2)^44^. Next, 3,003 cortical layer marker genes were identified by selecting the top 429 marker genes for each of the 6 cortical layers and the white matter from Table S4B of a recent publication^11^. Duplicated genes were removed, and the marker genes were intersected with our highly variable genes, resulting in 1,357 highly variable marker genes. The first 50 Principal Components (PCs) of these genes were calculated and normalized by Harmony (1.2.0) to remove batch effects^45^. Each section was considered a batch. Harmony was run with the parameters theta = 3, sigma = 0.1, and lambda = 1. Finally, BayesSpace (1.8.2) with spatial smoothing parameter gamma = 2 was applied to the first 35 harmony-normalized PCs to cluster spots into 7 clusters, corresponding to the 6 cortical layers and white matter^10^. White matter spots were not used in subsequent analyses.

Neuritic plaques were projected onto the ST coordinate system using the spot coordinates obtained from Space Ranger, and the distances between the plaques’ centers and the spots’ centers were calculated. Plaques were assigned to the cortical layer or white matter annotation that was most frequently observed at spots within a 200µm radius of the plaque, weighted by the inverse distance. GFAP MFI values per spot (Supplementary Figure 2) were calculated in ImageJ (1.54f) by mapping spot coordinates to raw images.

### Testing differential transcription at neuritic plaques

For the detection of up- and downregulated genes at neuritic plaques, only sections with at least one grey matter neuritic plaque were selected (24 sections from 14 donors). Spots were classified as plaque spot (≤150µm; n=1,448) or control spot (≥500µm; n=26,590) based on their distance to the nearest neuritic plaque. A mixed effects hurdle model, as implemented in the MAST package^46^, was fitted to the normalized spot-level data. The model included the spot status (plaque/control), the cortical layer, and the cell detection rate (CDR, fraction of genes detectable in the spot) as fixed effects, and the section as a random effect. Only genes detected in at least 5% of the spots were tested. The effect of the spot status variable was tested by a likelihood ratio test combining the effects from the logistic and Gaussian components of the hurdle model. Genes with an FDR less than 0.05 and an absolute log_2_ fold change (log_2_FC) larger than 0.05 were considered significant.

The explorative GO enrichment analysis for the biological process (BP) ontology was performed using the clusterProfiler package^47^. For the GO analysis, differential genes were defined using the same logFC cutoff but a less stringent FDR of 0.1 to obtain a larger set of differential genes. All tested genes were used as background.

### Deconvolution

Deconvolution ST data was performed using Cell2location (v0.1.3)^21^, a Bayesian model that integrates reference snRNA-seq data with ST data to resolve the cell states contributing to the transcriptomic signature of each ST spot. Reference snRNA-seq signatures were derived from the cell state annotations described in our previously published DLPFC cell atlas, which includes excitatory neurons (Exc), inhibitory neurons (Inh), astrocytes (Ast), oligodendrocytes (Oli), oligodendrocyte precursor cells (OPC), microglia (Mic), macrophages, and monocytes^2^. Microglial states representing less than 5% of total microglia (Mic.1, Mic.11, Mic.14, Mic.16), oligodendrocytic states representing less than 1% of total oligodendrocytes (Oli.10, Oli.11, Oli.12), and the rare neuronal state Inh.1 were excluded, resulting in a total of 67 cell types/states. While neuronal cells, oligodendrocytes, and OPCs were donor-matched, all nuclei identified as Ast and Mic from 424 donors in the reference dataset were used to increase the number of nuclei from less frequent glial cell types. Initial filtering was applied using the Cell2location parameters cell_count_cutoff=5, cell_percentage_cutoff=0.001, and nonz_mean_cutoff=1.06. The model was trained for 1,000 epochs (determined empirically).

After constructing the reference, we ran the Cell2location model with the hyperparameters N_cells_per_location=4, detection_alpha=10, and n_groups=50 for 8,000 iterations (determined empirically). Mitochondrial genes (HGNC gene group 1972), ribosomal genes, and mitochondrial ribosomal genes (HGNC gene group 1054) were removed from the ST data prior to cell state abundance estimation. We calculated the proportion of each cell state relative to its corresponding major cell type for each spot. Plaque and control spots were defined as in the differential transcription analysis, and the same 24 sections with at least one grey matter neuritic plaque were used. To test whether the proportion of a cell state differed between plaque and control spots, we logit-transformed the proportions and fitted a random-effects model for each cell state. The estimated proportion was used as the outcome, with plaque/control spot status, sex, and cortical layer as fixed effects, and section as a random effect, using the R package glmmTMB (1.1.9). The effect of plaque/control status was assessed using a Wald test and resulting p-values for all cell types and states were FDR-adjusted. To test for changes in cell state proportions between control spots distant from plaques in AD brains and spots from non-AD brains (Figure 3e), we removed all plaque spots and added our four control sections from non-AD brains. The same model was then applied, except that the plaque/control spot status was replaced with a variable encoding AD or control brain status.

### Immunohistochemistry staining and data analysis

#### Immunohistochemistry on FFPE tissue

6μm-thick sections of formalin-fixed paraffin-embedded DLPFC tissue obtained from the RUSH university were stained for GFAP-Cy3 conjugate (Millipore Sigma, MAB3402C3, dilution 1/100), SERPINA3 (Novus Biologicals, NBP1-90295, dilution 1/100) and AB_1-42_ biotinylated (Biolegend, Cat #805504, dilution 1/300). After deparaffinization using CitriSolv (DeconLabs, Cat#1601) for 20 min at RT and rehydration using ethanol washes, heat-induced epitope retrieval was performed using citrate (pH = 6; Sigma Aldrich, C9999) in a microwave for 25 min (400 Watts). The sections were treated with 88% formic acid (Fisher Scientific, A118P-100) for 2 min, blocked with 3% BSA (Sigma Aldrich, A7906) for 30 min, and incubated with the primary antibodies, anti-SERPINA3 and anti-AB_1-42_, overnight at 4°C. Sections were washed with PBS and incubated with fluorochrome-conjugated secondary antibodies for 1 hour at RT in the dark. The secondary antibodies used were donkey anti-rabbit Alexa Fluor 647 (Invitrogen, A31573), donkey anti-goat Alexa Fluor 488 (Invitrogen, A11055) and Streptavidin Alexa Fluor 730 (Invitrogen, S21384). After another round of blocking with 3% BSA for 30 min, the sections were incubated with GFAP-Cy3 conjugated primary antibody for 1 hour at RT in the dark. The sections were then washed again and incubated with True Black (Biotinum, Cat#23007) for 2 min at RT to quench Lipofuscin autofluorescence. They were then coverslipped with an anti-fading reagent with DAPI (Invitrogen, P36931), left at RT in the dark to dry, and stored at 4°C in the dark for subsequent imaging.

#### Imaging, astrocyte segmentation, and statistical analysis

Entire tissue sections were imaged using the Nikon Eclipse Ni-E epifluorescence microscope at 20X magnification (Plan Apo λ, NA = 0.75). Neuritic plaques were identified manually. Global coordinates were obtained by extracting image metadata with Bio-Formats (v5.7.1). Astrocyte objects were segmented using CellProfiler (v4.2.5)^48^ based on GFAP staining, and only GFAP^+^ objects overlapping with DAPI fluorescence were considered. Briefly, DAPI was segmented using the module ‘IdentifyPrimaryObjects’ with advanced settings. The diameter of objects was set to range between 18 and 80 pixels, and the ‘Robust Background’ method was used for thresholding. For GFAP segmentation, the ‘EnhanceOrSuppressFeatures’ module was used to enhance the ramification features of astrocytes. The ‘Line structures’ method was applied for enhancement. GFAP^+^ cells were then detected using the ‘IdentifyPrimaryObjects’ module, with the diameter of objects set to range between 10 and 300 pixels. ‘SplitOrMergeObjects’ was used to combine small objects, such as ramified structures, and assign them to nearby objects. GFAP^+^DAPI^-^ cells were filtered out using the ‘RelateObjects’ module, with DAPI defined as the parent objects and GFAP as the child objects. For each GFAP^+^DAPI^+^ object, the MFI of SERPINA3 and GFAP was measured.

The distance to the nearest plaque was measured for each GFAP^+^DAPI^+^ astrocyte. Astrocytes within 100μm of a plaque were classified as plaque-associated / proximal (n = 12,763 across 20 sections), while those 350–500μm away were considered control / distant astrocytes (n = 51,99 across 20 sections). The 500μm limit was set to avoid comparing plaque-associated astrocytes to distant astrocytes from different cortical layers, as cortical layer annotations were unavailable. Log_2_-transformed SERPINA3 MFI was used as the outcome variable in a mixed effects model, with astrocyte status (plaque-associated or control) and sex as fixed effects, and the nearest plaque nested within the section as random effects to account for variability between plaques and donors. The model was fitted using the R package glmmTMB (v1.1.9), and the effect of plaque association was assessed using a Wald test. The same model was used to test for increased GFAP MFI at plaques.

### DLPFC proteomic data analysis

Normalized tandem mass tags (TMT) proteomics data from the DLPFC of 595 ROSMAP participants were downloaded from Synapse (Synapse ID: syn51150434)^13^. Neocortical neuritic plaque burden was determined by microscopic examination of silver-stained slides from three neocortical regions (inferior parietal cortex, midfrontal cortex, and midtemporal cortex) and scaled and averaged to obtain a summary score. Neocortical neuritic plaque burden was available for a subset of 588 samples. SERPINA3 expression values were modeled as the outcome in a linear regression model with neuritic plaque burden as the explanatory variable and sex and age as covariates.

### CSF proteomic data analysis

The first CSF TMT proteomic dataset (Figure 2e) was downloaded from Synapse (SynID: syn20933797) and consisted of 150 healthy controls and 147 mild cognitive impairment/AD patients. Aβ_1-42_ was quantified in the CSF using an immunoassay as described in the original publication^14^ and was available for 296 out of the 297 samples. SERPINA3 expression values were modeled as the outcome in a linear regression model with Aβ_1-42_ as the explanatory variable, and sex and age as covariates.

The second CSF TMT dataset (Figure 2f) was accessed via the Alzheimer’s Disease Data Initiative (ADDI) (https://doi.org/10.58085/HR6S-2991)^15^. Since quantified Aβ_1-42_ measures were not available, we compared 187 control samples (normal cognition and normal CSF biomarker profile) to 209 samples from patients with AD dementia diagnosis confirmed by CSF biomarkers, using the normalized SERPINA3 levels as the outcome of a regression model adjusted for sex and age.

## Data Availability

The raw and processed data generated by this study (spatial transcriptomic and imaging data) are available via the AD Knowledge Portal (https://adknowledgeportal.org). The AD Knowledge Portal is a platform for accessing data, analyses, and tools generated by the Accelerating Medicines Partnership (AMP-AD) Target Discovery Program and other National Institute on Aging (NIA)-supported programs to enable open-science practices and accelerate translational learning. The data, analyses and tools are shared early in the research cycle without a publication embargo on secondary use. Data is available for general research use according to the following requirements for data access and data attribution (https://adknowledgeportal.org/DataAccess/Instructions).

For access to content described in this manuscript see: https://doi.org/10.7303/syn62110225 Pathologic and phenotypic data from ROSMAP are available on our Resource Sharing Hub at https://www.radc.rush.edu.

## Author information

*These authors contributed equally: Mariko Taga, Hans-Ulrich Klein

### Authors and Affiliations

**Center for Translational & Computational Neuroimmunology, Department of Neurology, Columbia University Irving Medical Center, New York, NY, USA**

Berke Karaahmet, Ya Zhang, Laurine Duquesne, Alina Sigalov, Christina Yung, Alexandra Kroshilina, Mariko Taga, Hans-Ulrich Klein

**Rush Alzheimer’s Disease Center, Rush University Medical Center, Chicago, IL, USA**

David A. Bennett

## Contributions

MT and HK conceived and designed the study. YZ, LD, AS, CY, AK, and MT conducted the combined ST and IF experiments. BK, AS, and MT performed the IHC validation experiments. BK, MT, and HK analyzed the data. BK and HK drafted the manuscript. All authors interpreted the results and approved the final version of the manuscript.

### Corresponding Authors

Correspondence to Hans-Ulrich Klein or Mariko Taga

## Acknowledgements

This work was supported by an Alzheimer’s Association Grant through the AD Strategic Fund (ADSF-21-816675). The ROSMAP studies were supported by grants from the National Institutes of Health (P30AG10161, P30AG72975, R01AG15819, R01AG17917, U01AG46152, and U01AG61356).

