## Supplementary Figures for "Spatial Transcriptomic Analysis Identifies a *SERPINA3*-Expressing Astrocytic State Associated with the Human Neuritic Plaque Microenvironment"

**This .pdf file includes:**

Supplementary Figures 1 – 5
Supplementary Table Captions
References

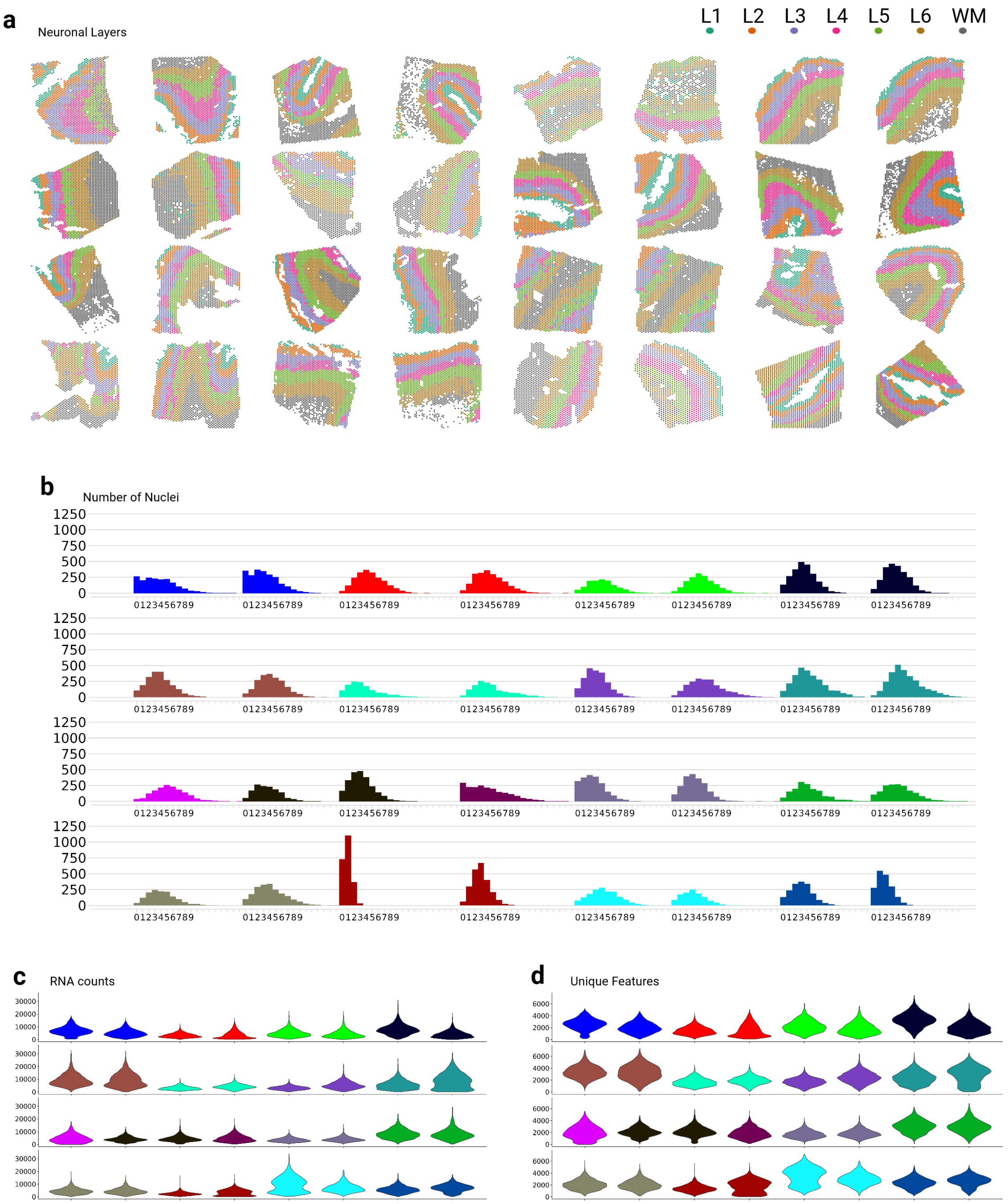


**Supplementary Figure 1.** General characteristics of the ST dataset. **(a)** Layer annotations showing the anatomical structure of the 32 DLPFC tissue sections characterized in this study. **(b)** Histograms of the number of DAPI⁺ nuclei per spot. Different colors depict different tissue donors. **(c-d)** Violin plots of RNA counts **(c)** and unique features **(d)** per tissue section.

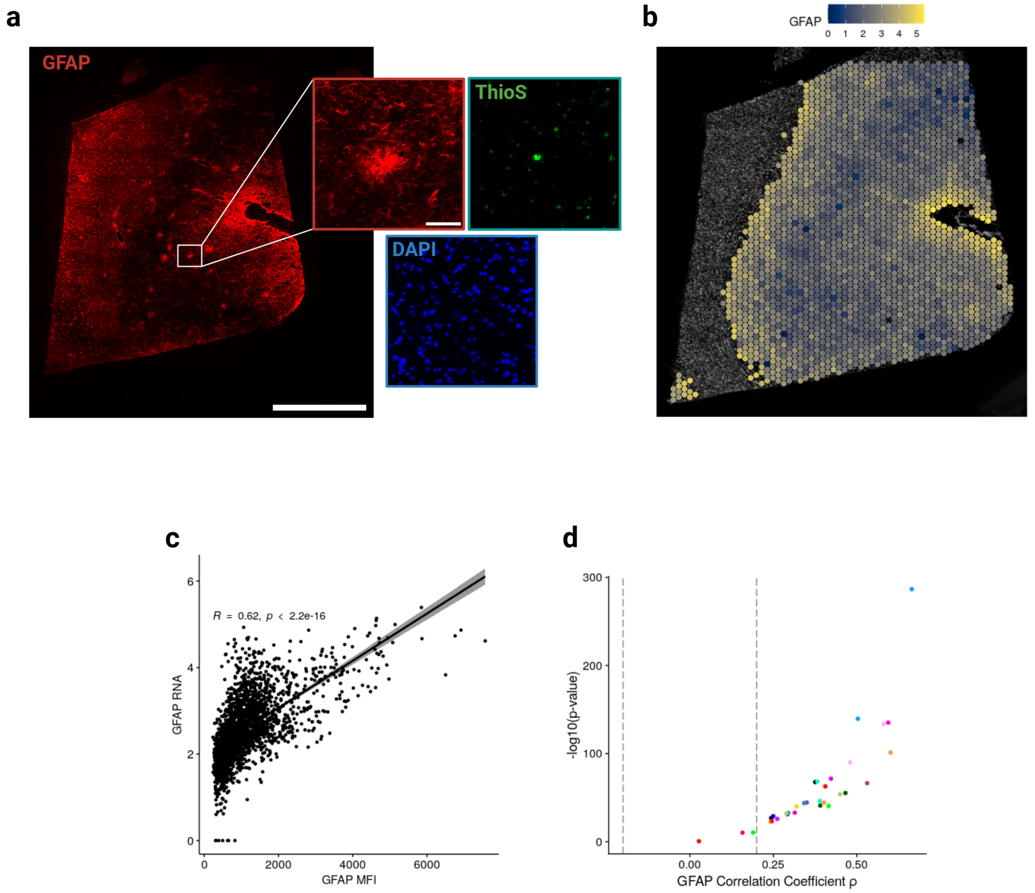


**Supplementary Figure 2.** Normalized *GFAP* counts from ST spots correlate with GFAP MFI from the IHC images. **(a)** Representative IHC staining showing GFAP (red), ThioS$^{+}$ neuritic plaques (green) and DAPI (blue). Scale bars denote 500µm and 25µm in their respective magnification orders. **(b)** Representative ST section of the same tissue shown in (a) illustrating normalized *GFAP* mRNA counts. **(c)** Correlation between GFAP MFI from (a) and *GFAP* RNA from (b) for each ST spot. **(d)** Correlation coefficients and their p-values are shown for all 32 tissue sections in the ST dataset. Dashed lines represent Spearman's correlation coefficient (ρ) of |0.2| and colors denote donors. A significant positive correlation was observed for all sections except one.

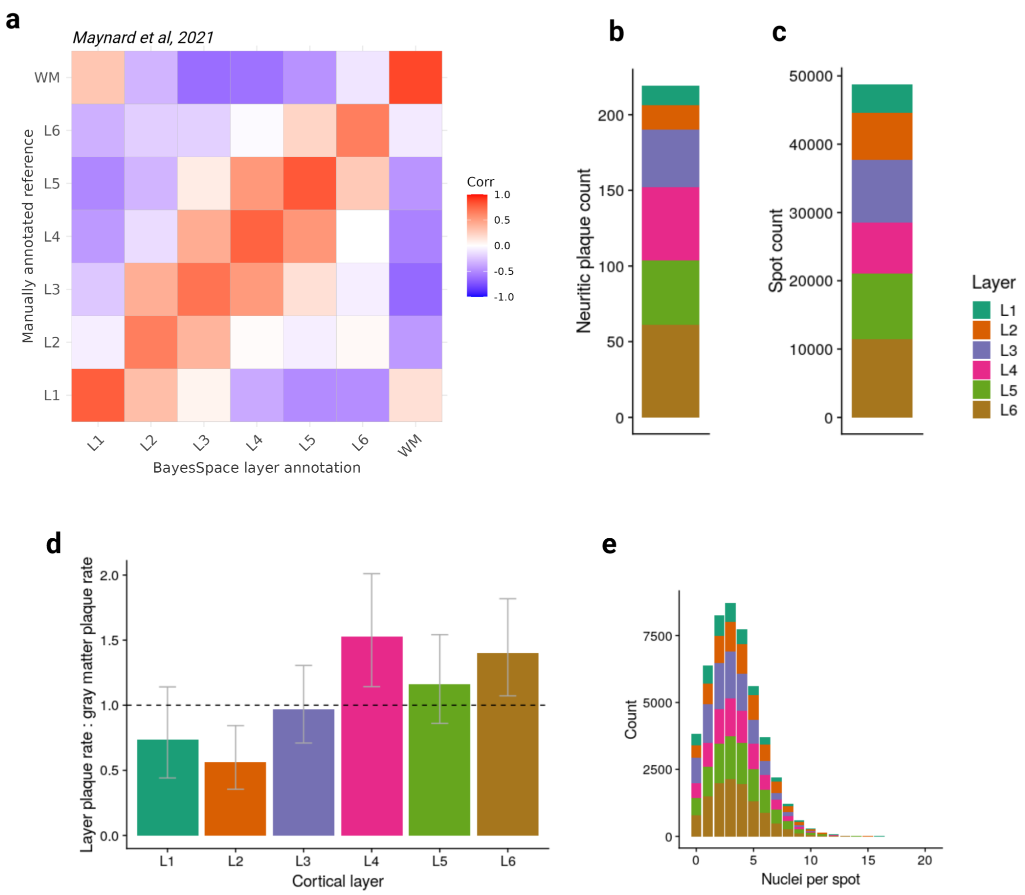


**Supplementary Figure 3.** Characteristics of the computationally identified neuronal layers. **(a)** Applying BayesSpace algorithm to computationally annotate layers correlates strongly with the manual H&E-based layer annotation of ST spots in the Maynard et al. ST dataset. First, pseudobulk data were generated by summing the counts for each layer per sample, followed by TMM normalization. Differential expression testing was then performed by comparing each individual layer to all other layers, yielding a vector of t-statistics that describe gene up- and downregulation in the respective layer. This procedure was applied to our dataset, annotated with BayesSpace (x-axis), and to the dataset from Maynard et al., which was manually annotated (y-axis). The heatmap displays the Pearson correlation between the t-statistic vectors obtained from the two datasets. **(b-c)** Number of neuritic plaques **(b)** and number of spots **(c)** per layer in our ST dataset. **(d)** Plots showing enrichment of neuritic plaques in layers 4-6, compared to other layers. Bars indicate the layer-specific plaque rate divided by the mean gray matter plaque rate. Error bars show

95% confidence intervals. The plaque rates differed significantly between layers in our dataset (p=0.001, Poisson regression model with donor as random effect) **(e)** Histogram showing the distribution of nuclei per spot by layer, with most spots containing 4 nuclei.

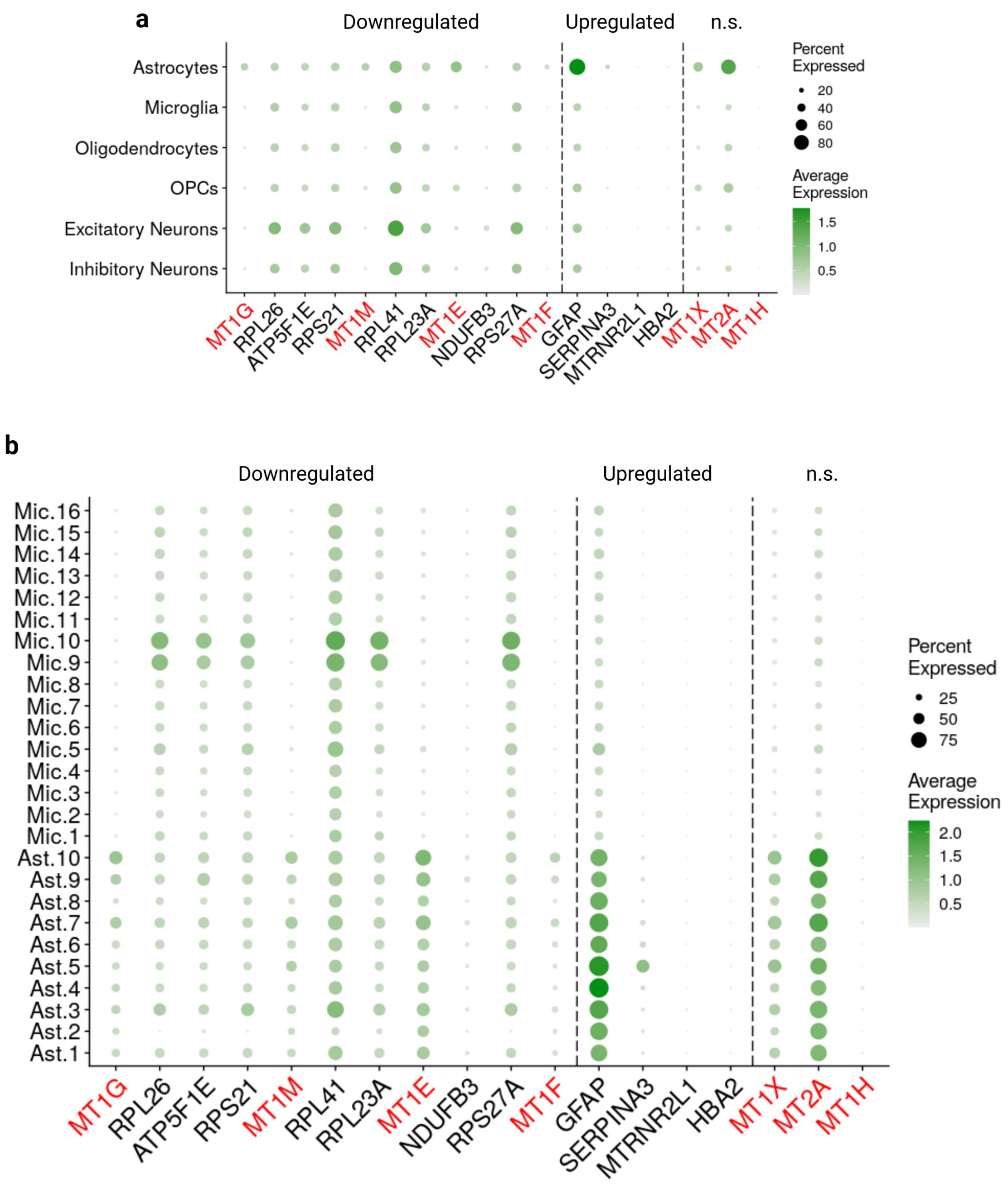


**Supplementary Figure 4.** Expression of neuritic plaque DEGs in the Green et al. snRNA-seq dataset**. (a)** SCTransformed expression of the top 10 downregulated and significantly upregulated genes identified in our ST dataset in six major cell types of the brain. Additional metallothionein genes detected in the ST dataset, though not reaching significance, were also included. *SERPINA3* expression was limited to astrocytes. Although metallothioneins (red) were broadly expressed, they appeared to be most prominently expressed by astrocytes. **(b)** Microglial and astrocytic state-level expression of those genes. High levels of *GFAP* and *SERPINA3* expression were particularly noticeable in Ast.5. n.s., not significant.

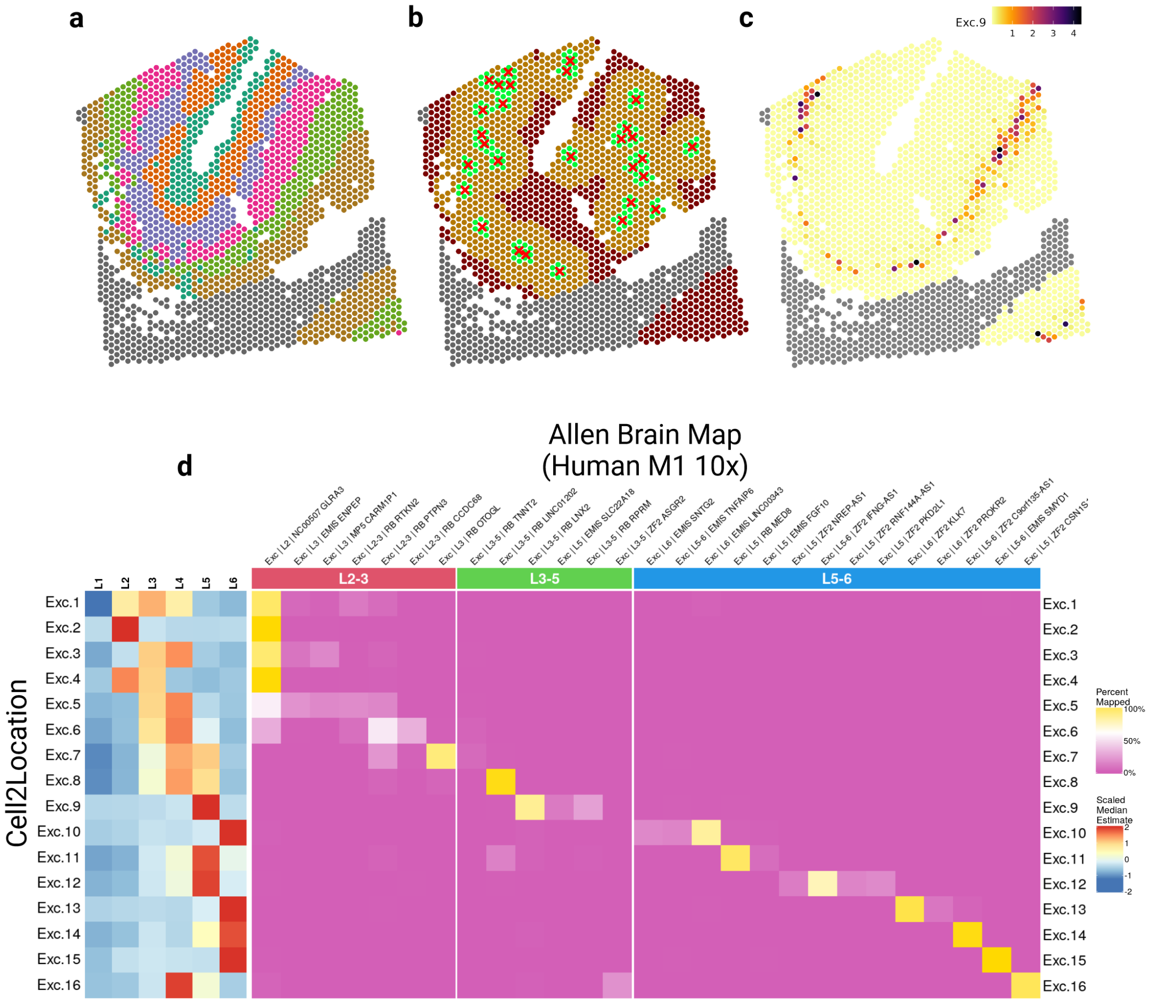


**Supplementary Figure 5.** Validation of computational neuronal layer annotation using spatial deconvolution. **(a)** A representative tissue section and its neuronal layer annotation. **(b)** Illustration of the neuritic plaque distribution (green < 150µm, orange < 500µm) and **(c)** the estimated abundance of excitatory neuronal state 9 (Exc.9) in that tissue section. **(d)** Heatmaps showing the enrichment of various deconvoluted excitatory neuronal states in ST spots assigned to given neuronal layers annotated through BayesSpace clustering (left), or label transfer of excitatory neuronal states from the reference snRNA-seq dataset to Allen Brain Map neuronal layer annotations (right). The left heatmap is derived from ST data and its deconvolution, while the heatmap on the right uses snRNA-seq label-transfer between Green et al. and Allen Brain Map (Human-M1-10X). The mapping of excitatory neuronal states to anatomical layers is similar between cell2location output with Green et al. reference snRNA-seq and label-transfer to a different, manually annotated dataset.

**Supplementary Table 1.** Demographics of the DLPFC tissue donors for the ST dataset and the IHC dataset. “NIA Reagan Diagnosis” is the NIA-Reagan diagnosis of AD which is based on neurofibrillary tangles (braaksc) and neuritic plaques (CERADsc)[1]. “CERAD Score” is a semiquantitative measure of neuritic plaques as recommended by the Consortium to Establish a Registry for Alzheimer's Disease (CERAD)[2].

**Supplementary Table 2.** Characteristics of the Visium dataset, showing the number of ST spots and neuritic plaques per layer for each tissue section.

**Supplementary Table 3.** Output of the differential expression testing of ST spots within 150μm of a neuritic plaque versus spots further than 500μm away from the closest plaque.
